# AN ACCURATE GENETIC CLOCK

**DOI:** 10.1101/019414

**Authors:** D. H. Hamilton

## Abstract

Molecular clocks give *“Time to most recent common ancestor” TMRCA* of genetic trees. By Watson-Galton^17^ most lineages terminate, with a few overrepresented *singular lineages* generated by W. Hamilton’s “kin selection”^13^. Applying current methods to this non-uniform branching produces greatly exaggerated *TMRCA.* We introduce an inhomogenous stochastic process which detects *singular lineages* by asymmetries, whose *reduction* gives true *TMRCA.* This implies a new method for computing mutation rates. Despite low rates similar to mitosis data, *reduction* implies younger *TMRCA,* with smaller errors. We establish accuracy by a comparison across a wide range of time, indeed this is only clock giving consistent results for both short and long term times. In particular we show that the dominant European y-haplotypes R1a1a & R1b1a2, expand from c3700BC, not reaching Anatolia before c3300BC. While this contradicts current clocks which date R1b1a2 to either the Neolithic Near East^4^ or Paleo-Europe^20^, our dates support recent genetic analysis of ancient skeletons by Reich^23^.

---

The genetic clock, computing *TMRCA* by measuring genetic mutations, was conceived by Emile Zuckerkandl and Linus Pauling ^32^,^33^ on empirical grounds. However work on neutral mutations by Motto Kimura^16^ gave a theoretical basis and formula. While our theory applies to general molecular evolution, we focus on the Y-chromosome with DYS regions (**D**NA **Y**-chromosome **S**egments) counting the “short tandem repeat” (STR) number of nucleotides of a micro satellite. In fact one uses many DYS sites, marked by *j* = 1, *…N*, each individual *i,* 1 = 1, *‥n,* has STR number *x*_*i,j*_. The Y-chromosome is passed unchanged from father to son, except for mutations *x*_*i,j*_ → *x*_*i,j*_ *±* 1 occurring at rate

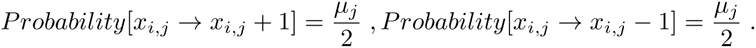

The fundamental assumption is that the sample population has a single patriarch at time *t* = *TMRCA*(generations). Now suppose the present (sample) population has mode *m*_*j*_ at DYS *j*. This is taken to be the STR value of the original patriarch. A calculation shows the present population with variance 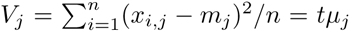. Then averaging over the markers gives *TMRCA* = Σ_*j*_ *V*_*j*_ / (*n* Σ_*j*_ *μ*_*j*_). This variance method and its variations we call KAPZ after its originators.

In practise problems soon arose. Mutation rates could be computed from mitosis, but sample sizes are too small to give great accuracy. Using these a KAPZ due to Zhivotovsky^31^,^31^ was applied to R1b1a2 by Myres^20^ giving L23*(Turkey) giving 9000BC, *σ* = 2000.

Mutation rates could also be estimated from large family groups with genealogy data. However there are significant discrepancies in rates between different family groups. Also these “pedigree” rates are much larger than those from mitosis. A similar phenomena for the mitochondrial clock suggested high short term rates and lower long term rates^14^,^15^. So very low long term rates of .00069 were suggested^31^ for the Y-clock. We show this is unnecessary.

Another problem is that KAPZ is for large populations whereas ancient populations were small and modern samples can be tiny, e.g. *n <* 20. This led to the introduction of Bayesian methods such as BATWING^27^, which considers all possible genealogical trees giving the present sample data, then searches for the tree of maximum likehood. But the BATWING *TMRCA* is often greater than KAPZ, e.g. for the Cinnioglu^8^ study of Anatolian DNA both methods were applied to the same data and mutation rates. For R1b1a2 the KAPZ had *T M RCA* = 9800*BC* compared with 18, 000*BC* for BATWING. Balaresque^4^ used BATWING to give an origin for R1b1a2 in Neolithic Anatolia c. 6000BC, but their statistics was disputed by Bushby^29^. All of this was contradicted by Reich^22^ who found R1b1a2 in skeletons c 3300BC from Yamnaya cemeteries.

**Figure 1:**
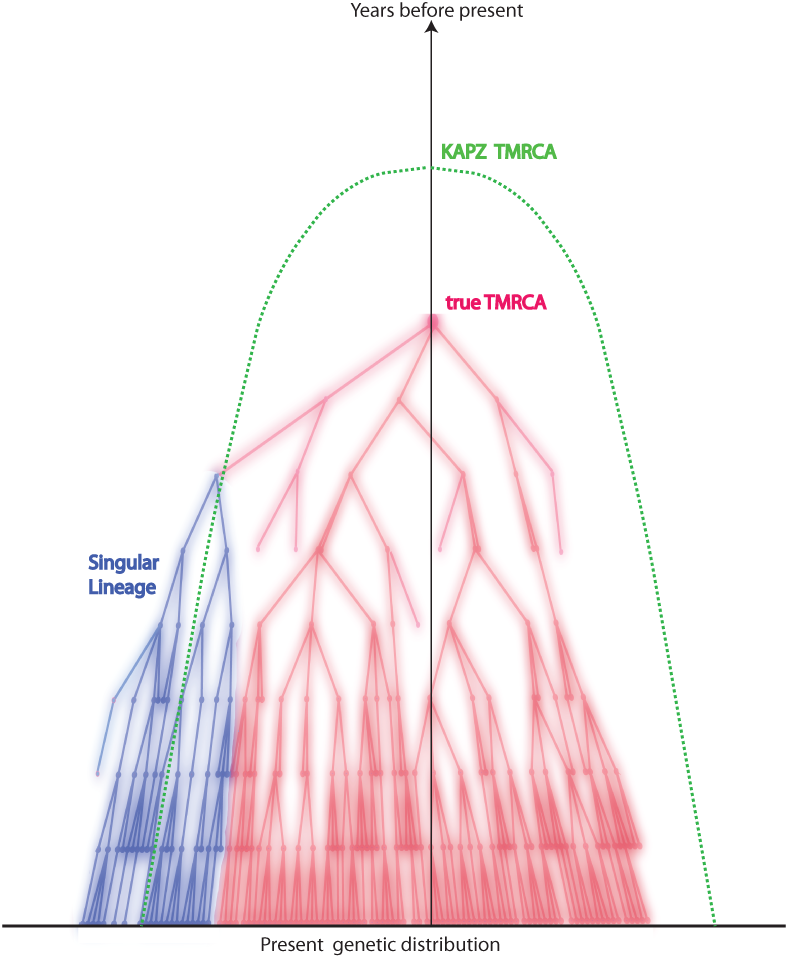
A singular lineage increases variance and apparent TMRCA:

## Singular Lineages

A fundamental problem is that present populations have highly overrepresented branches we call *singular lineages*. A well known example is the SNP L21 which is a branch of R1b1a2. Individuals identified as L21 are often excluded from R1b1a2 analysis because they skew the results. Such a singular lineage causes the variance to be much greater, even though the original *TMRCA* remains unchanged, see figure 1. For Bayesian methods such lineages are very unlikely giving an even greater apparent *TMRCA*. However one cannot deal with singular branches by excluding them. For one thing, our method will show that 50% of DYS show evidence of singular side branches, i.e. more than a SD from expected. Excluding them would also remove some of the oldest branches and produce a *TMRCA* which is too young. Now these singular lineages are very (mathematically) unlikely to arise from the stochastic system which is the mathematical basis of KAPZ (or the equivalent Monte-Carlo process modeling BATWING). We believe that the standard stochastic process is perturbed by other improbable events, which are then amplified by biological processes.

First, the Watson-Galton Process^17^ implies lineages almost certainly die out. Conversely, the “kin selection” of W.D. Hamilton^13^, shows kin co-operation gives genetic advantages. Consider three examples with well developed DNA projects. Group A of the Hamiltons has approximately 100, 000 descended from a Walter Fitzgilbert c 1300AD. Group A of the Macdonalds has about 700, 000 descendants from Somerfeld c1100AD, and Group A of the O’Niall has over 6 million descendants from Niall of the Seven Hostages, c300AD. These are elite groups with all the social advantages. One sees lines of chieftains, often polygamous. Our model has many extinct twigs with a few successful branches, whereas current models assume a uniform “star radiation”, see below

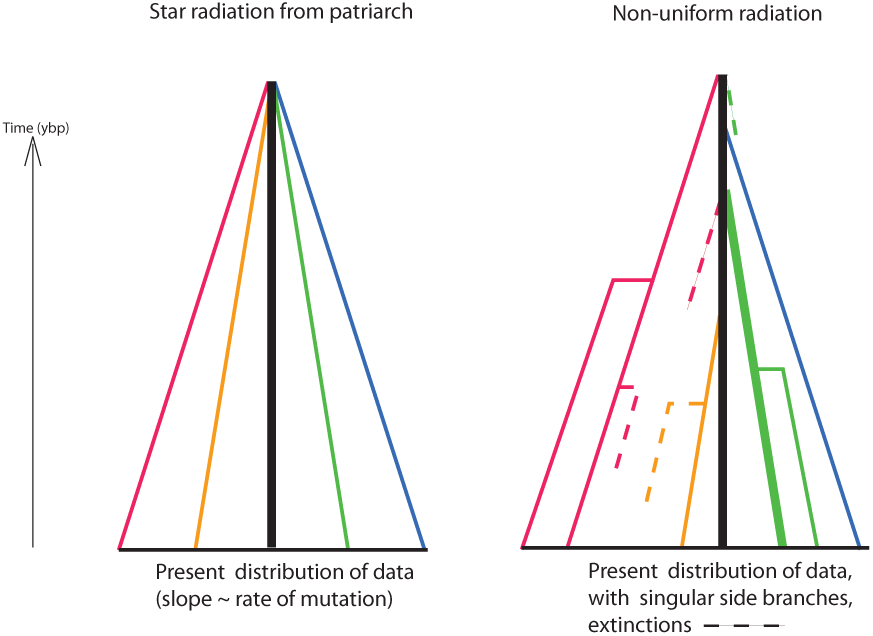

## Reduction of Singular Lineages

Modelling singular lineages requires a new stochastic system where instead of a single patriarch we imagine many “virtual patriarchs”, each originating at tme *t*_*k*_ ago. Each of these giving a proportion 0 ≤*ρ*_*k*_ ≤1 of the present population. So we now have an inhomogenous expansion. Furthermore the symmetric model for mutations has to be changed to

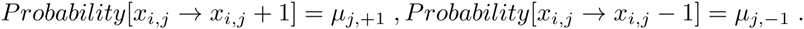

We introduce asymmetric mutations and show how to compute it. Asymmetry will play a very important role in detecting singular lineages. This inhomogenous asymmetric system is mathematically equivalent to a mixed population. Computing its solution is an “inverse problem”. Unfortunately inversion is unstable for such systems, also there is no unique solution. However it turns out that, up to a standard deviation SD, most DYS markers show at most one singular branch which is found from asymmetries in the distribution. These singular branches are then *reduced* revealing the original lineage. We then compute a branching time *t*_*j*_ for each marker *j*. The effect of reduction is dramatic, see Figure 3. Now the nonuniform branching process causes the *t*_*j*_ to be ran-domly distributed so their mean is not the TMRCA. Large errors in mutation rates means one cannot simply take the max *t*_*j*_ to be the *TMRCA*. Instead stochastic simulations of the branching process, using robust statistics to avoid outliers, find the most likely *TMRCA*, see Supplementary Material 1 (SM1) for full mathematical details.

These methods also imply a new way of computing mutation rates, see SM2. Previously, there were methods based on mitosis data or pedigree studies of family DNA projects (which gave quite different rates). We begin with 8 very large SNP projects from FTDNA using 37 markers, of course with unknown *TMRCA* and find mutation rates as the fixed points of a stochastic process. These take about 3 iterates to converge. After we discard markers with mutation *SD >* 33% we are left with 29 markers. We find the mutation rates are close to those obtained from mitosis and nearly 1/3 the values obtained by pedigree. Despite the fact that our mutation rates are lower than most studies, *reduction of singular lineages* produces more recent *TMRCA* than current models.

## Examples

Our clock is the only one with across the board consistent results:

**Table 1:**
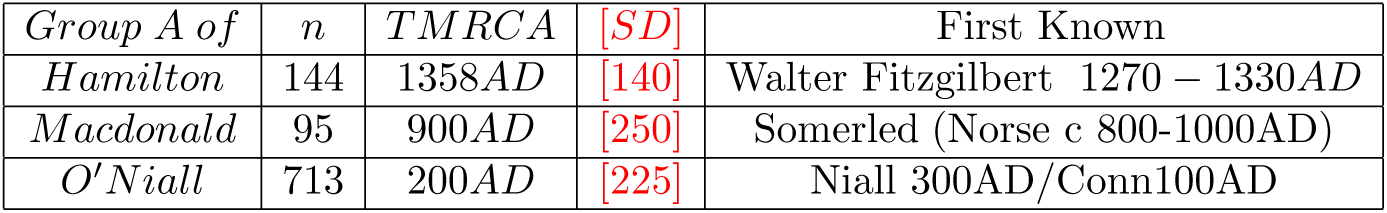
*TMRCA* for Medieval groups.

Archeological finds convinced Marija Gimbutas^11^ to attribute Proto Indo-European (PIE) to the Yamnaya Culture c 3500BC of the Russian Steppes, see Anthony^2^. This is consistent with mainstream linguistic theory, some even wrote of linguistic DNA. But actual genetics was ignored because this contradicts current genetic clocks. Now the dominant European y-haplotypes are R1b1a2 & R1a1a (which like other y-haplotypes is marked by a unique single nucleotide polymorphism (SNP) mutation). Table 2 shows the expansion times of c3700BC, similar for regions Russia, Poland, Germany and Scandinavia. The times are so close only Scandinavia is significantly later. This data is from FTDNA projects for region X only using individuals with named ancestor from These independent results agree within the standard deviation (SD), with dates matching the Corded Ware Culture, a semi-nomadic people with wagons and horses who expanded west from the Urkraine *c*3000*BC*. This is consistent with the oldest R1b1a2, R1a1a skeletons being from the Yamnaya Culture^23^.

**Table 2:** R1b1a2, R1a1a independent comparison

| *Region* | *R1b1a2* | *[SD]* | *n* | *R1a1a* | *[SD]* | *n* |
| --- | --- | --- | --- | --- | --- | --- |
| *All* | 3700BC | [625] | 460 | 3800BC | [700] | 1270 |
| *Russia* | NA |  |  | 3750BC | [700] | 337 |
| *Poland* | 3960BC | [950] | 65 | 4600BC | [820] | 876 |
| *Germany* | 2780BC | [500] | 438 | 3750BC | [800] | 190 |
| *Scandinavia* | 2550BC | [500] | 153 | 4500BC | [1000] | 140 |

An interesting intermediate step occurs between the medieval and eneolithic. The mythical Irish Chronicles relate that the O’Niall descend directly from the first Gaelic High Kings, which tradition dated c1300-1600BC. The O’Niall have the unique mutation M222 which is a branch of the haplotype L21. For L21, *n* = 1029, we compute *TMRCA* = 1600*BC* and SD *σ* = 320. These are dates for proto Celtic, i.e. what archeologists call the pre Urnfelder Cultures, c. 1300-1600BC, see SM5. Furthermore L21 is in turn a branch of haplotype P312 which we date to 2300BC. This date suggests the Bell Beaker Culture of Western Europe. Indeed the only known^23^ Bell Beaker genome is P312 with ^14^*C* date 2300BC.

Our method requires large data sets and many markers which means we have to rely on data from FTDNA, finding 29 useable markers out of standard 37 they use. In fact many researchers^4^ have used FTDNA data. We think our method of reduction with robust statistics solves any problems with this data. To test this we compared our results with R1a1a1 data obtained from Underhill^26^ with *n* = 974(which involved excluding his four M420 individuals and others with missing markers), and 15 useable markers. The result was 2550*BC, σ* = 400, within the CI of our R1a1a results. Table 5 shows the results of extensive simulations using random subsets of our FTDNA data, for 29, 15 and 7 markers. For the same 15 markers as the Underhill^26^ the different FTDNA data gives very similar 3300*BC, σ* = 840 for R1a1a, verifying the correctness of using FTDNA data. However once you get down to 7 markers the confidence interval becomes large, e.g. R1a1a gives 3400*BC, σ* = 1500. Also it becomes difficult to deal with outliers.

An example with few markers is R1b1a2 data of Balaresque^4^. Our method (this time with 7 useable markers) gave SD *>* 30%, see Table 6. Now Balaresque^4^ used the Bayesian method BATWING^29^ to suggest a Neolithic origin in Anatolia. With the same Cinnioglu^8^ data our method gives for Turkish R1b1a2 (*n* = 75) a *TMRCA* = 5300*BC, σ* = 3100, i.e. anytime from the Ice Age to the Iron Age. Fortunately, once again, we find good data from FTDNA: the Armenian DNA project, see Table 3. By tradition the Armenians entered Anatolia from the Balkans *c*1000*BC* so they might not seem a good example of ancient Anatolian DNA. But some 100 generations of genetic diffusion has resulted in an Armenian distribution of Haplotypes J, G, R1b1a2 closely matching that of all Anatolians, therefore representive of typical Anatolian DNA. We see that Anatolian R1b1a2 arrived after c3300BC, ruling out the Neolithic expansion c6000BC. When dealing with regional haplotypes, e.g. R1b1a2 in Anatolia, the *TMRCA* is only a upper bound for the arrival times, for the genetic spread may be carried by movements of whole peoples from some other region.

| Armenian | $n$ | $TMRC A$ | $[SD]$ |
| --- | --- | --- | --- |
| R1b1a2 | 99 | 3300BC | [800] |
| G2a2b | 46 | 9300BC | [2000] |
| J2 | 97 | 12100BC | [2200] |

Observe that our *TMRCA* for Armenian G2a2b (formerly G2a3) and J2 show them to be the first Neolithic farmers from Anatolia, i.e. older than 7000*BC*. In Table 4 we compared J2, G2a2b for all of Western Europe (non-Armenian data). Our dates show J2 was expanding at the end of the Ice Age. Modern J2 is still concentrated in the fertile crescent, but also in disconnected regions across the Mediterranean. The old genetic model predicted a continuous wave of Neolithic farmers settling Europe. But you cannot have a continuous maritime settlement: it must be *leap-frog*. Also repeated resettlement from the Eastern Mediterranean has mixed ancient J2 populations, and our method gives the oldest date. On the other hand G2a2b shows exactly the dates expected from a continuous wave of Neolithic farmers across Central Europe, consistent with Neolithic skeletons showing G2a2b (e.g. the famous Iceman).

| $SNP(n)$ | 7 marker | $[SD]$ | 15 marker | $[SD]$ | 29 marker | $[SD]$ |
| --- | --- | --- | --- | --- | --- | --- |
| G2a2b(1221) | 4800BC | [2050] | 8600BC | [2120] | 5359BC | [900] |
| R1b1a2(460) | 5524BC | [2000] | 4300BC | [950] | 3700BC | [625] |
| R1a1a(1270) | 3400BC | [1500] | 3200BC | [840] | 3800BC | [700] |
| I1(2898) | 3500BC | [1500] | 2711BC | [950] | 1800BC | [400] |
| L21(1029) | 1870BC | [800] | 1700BC | [400] | 1600BC | [325] |
| U106(1533) | 1800BC | [800] | 2500BC | [600] | 2400BC | [440] |
| J2(1241) | 6100BC | [2100] | 18500BC | [3000] | 15500BC | [2600] |
| P312(971) | 2600BC | [900] | 2850BC | [625] | 2240BC | [420] |

## Discussion

History, archeology, evolutionary biology, not to mention epidemics (e.g. dating HIV), forensic criminology and genealogy are just some of the applications of molecular clocks. Unfortunately current clocks have been found to give only “ballpark” estimates. Our method is the only one giving accurate time, at least for the human y-chromosome verified over the period 500 - 15, 000*ybp*. Our methods should also give accurate times for mitochondrial and other clocks.

Many geneticists thought natural selection makes mutation rates too variable to be useful. The problem is confusion between the actual biochemical process giving mutations and superimposed processes like kin selection producing apparently greater rates. Notice that the SD for our mutation rates is on average 14% which is much smaller than the actual previous rates. We believe this small SD proves the reality of neutral mutation rates of Moto Kimura^16^.

While our method is accurate for “big data”, applications to genetics, forensics, genealogy require the *TMRCA* between just two individuals, or between two species. Now for this “2-body problem” we cannot determine what singular lineages the branching has been through: with mutations either exaggerated or suppressed. Thus previous methods for small samples are at best unreliable. It is an important problem to find what accuracy is possible for small samples.

In checking accuracy we ran into the question of the origins of PIE. Although there are genes for language there is certainly none for any Indo-European language. Thus inferences have to be indirect. Marija Gimbutas saw patterns in symbolism and burial rituals suggesting the Yamnaya Culture was the cradle of Proto Indo-European. Also their physiology was robustly Europeanoid unlike the gracile skeletons of Neolithic Europe, but this could be nutrition and not genetic. So it was an open question whether the spread of this robust type into Western Europe in the late Neolithic marked an influx of Steppe nomads or a revolution in diet.

Reich^23^ observed all 6 skeletons from Yamnaya sites, c 3300BC by ^14^*C* dating, are either R1a1b1 and R1a1a. But that method could not date the origin of R1a1b1 and R1a1a. Our *TMRCA* shows both these haplotypes expanding at essentially the same time c3700BC. This, together with our later date for Anatolia, implies that R1b1a2 and R1a1a must have originated in the Yamnaya Culture, c 3700BC. Furthermore, considering the correlation of haplotypes R1b1a2 and R1a1a with Indo-European languages (i.e. all countries with R1b1a2 & R1a1a frequency *>* 50% speak Indo-European), this provides powerful evidence for the origin of Proto Indo-European.

## Supplementary Material 1: Mathematical Genetics

Sophisticated Mathematical theory has been developed for genetics, see ^1,2,5^. However all of these assume the present distribution is entirely the result of the stochastic process. We emphasize the role of extraneous forces like kin-selection which operates on too big a scale and rarely enough with results that cannot be subsumed into the mutation rates. So our method does not follow from any of these previous theories nor is it just applying a known statistics package. Instead we return to basic principles.

### Fundamental Solutions

The Y-chromosome has DYS marked by *j* = 1, *…N*, where one can count the STR number *x*_*j*_. Consider the probability *P*_*j,k*_ (at time *t* generations) that at marker *j* we have *x*_*j*_ = *k*. This satisfies the homogenous stochastic system

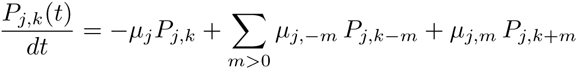

This homogenous system gives a uniform expansion from a single patriarch.

The system is essentially the model of Wehrhahn^9^ who had *μ*_*j,-*1_ = *μ*_*j,*1_. We introduce asymmetric mutations with total rate

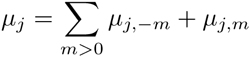

About 50% of DYS markers show asymmetric mutations, i.e. *μ*_*j,-*1_ *≠ μ*_*j,*1_.

The fundamental solution comes from the generator function

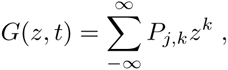

with complex variable *z*, and normalized initial condition *x*_*j*_ = 0 or *P*_*j,*0_(0) = 1:

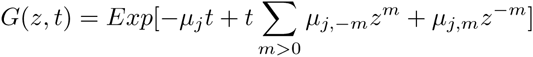

Then *G* can be expanded in powers of *z* to give *P*_*j,k*_(*t*). Now for the simplest asymmetric case, with only one step mutations, we have *G*(*z, t*) =

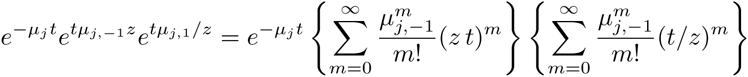

so using the Hyperbolic Bessel Function of Order *k ≥* 0, see Olver ^7^

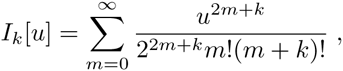

we see that the homogenous system has fundamental solution

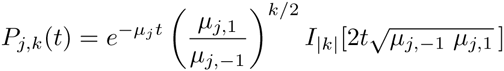

From this we obtain the second moment:

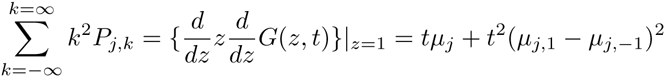

Also from the fundamental solution we find, independently of time

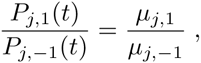

which we call the *asymmetric ratio*. It will be repeatedly used.

Of course the actual initial value is not *x*_*j*_ = 0 but was usually taken to be the mode *m*_*j*_ which was assumed to be the value for original patriarch. Assuming symmetry, i.e. *μ*_*j,-*1_ = *μ*_*j,*1_, the TMRCA is:

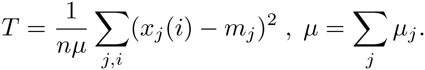

From the present distribution of data we use the frequency

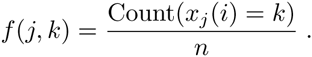

One problem with the KAPZ formula is that higher frequencies *f* (*j, k*), *|k|* = 2, 3*…* are overrepresented in the actual data. This is because the probability of a spontaneous two step mutation is much higher then the product of two one step mutations. So instead we use the frequency to solve the transcendental equation for the unknown *t*

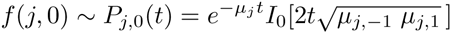

This nonlinear equation is easily solved via mathematical software such as MATHEMATICA (I used version 9 running on a boosted 2014 iMac which has accurate hyperbolic Bessel functions. Earlier versions on older iMacs gave inaccuracies so one had to compile one’s own functions). Using this formula resolves some other problems with the KAPZ method, e.g. *μ*_*j,-*1_ ≠ *μ*_*j,*1_ gives an extra quadratic term which if ignored causes large errors.

### Heterogeneous diffusion equation

However the main problem is singularities in the stochastic process. For a uniform stochastic process, 1 - *P*_*j*,0_(*t*) *∼* 1 *- f* (*j,* 0) is the probability of some mutation. So the expected variance is *f* (*j,* 0)(1*-f* (*j,* 0)). Thus if the actual data variance *V*_*j*_ *>> f* (*j,* 0)(1 *-f* (*j,* 0)) we are not uniform. Now a sublineage of very high fertility increases variance, giving apparently greater *TMRCA* although it is unchanged. One finds similar results for Bayesian methods.

The correct approach to nonuniformity assumes at times *t*_*i*_ (generations ago) a certain proportion 0 *≤ ρ*_*i*_ *≤* 1 of the present population originated from a “virtual patriarch” with an initial STR value *m*_*i*_. The resulting system :

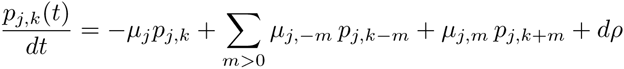

i.e. *dρ* are atoms of weight *ρ*_*i*_ with STR value *m*_*i*_ occurring at time *t*_*i*_. As the system is linear and isotropic the solution is a combination of fundamental solutions *P* of the homogenous system. Thus the present distribution *f* (*j, k*) is

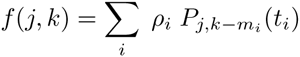

This allows us to consider populations mixed by having singular lineages from overfertile patriarchs, or by actual immigration from the outside. The inverse problem seeks to find singularities from present data. Unfortunately inversion is ill posed for such systems like the heat equation. This instability produces poor accuracy. Furthermore there is no unique solution, e.g.the present distribution could have been created yesterday.

However we find that *∼* 50% of the DYS markers show no significant difference from the uniform expansion of a single patriarch, i.e. the data variance *V*_*j*_ is close to the expected variance *f* (*j,* 0)(1 *-f* (*j,* 0)). The other markers show at most one significant side branch, i.e. there is an original branch starting at time *t*_*j,*0_ with STR *m*_0_ and a second one with STR *m*_1_ = *m*_0_ *±* 1 at time *t*_*j,*1_ *< t*_*j,*0_ with significant 0 *< ρ*_1_ *< ρ*_0_.

### Reduction

We locate these singular lineages by looking for asymmetries in the distribution. For a uniform flow from a single patriarch the frequency of STR value *k* is given by *f* (*j, k*) *∼ P*_*j,k*_(*t*). The asymmetric ratio:

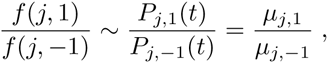

is completely independent of time *t*. Therefore if say

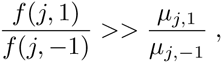

we have a singular lineage at *k* = +1. Thus the excess at *k* = +1 is

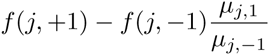

To first order approximation then frequency *f* (*j,* +2) is due to this singularity at *j* = +1 which therefore gave a contribution

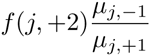

to *k* = 0. Thus removing the effect of the singularity at *k* = +1 leads to new frequencies

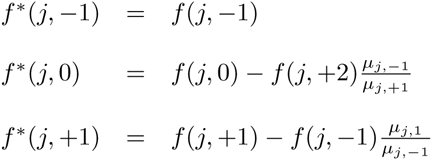

These of course are no longer normalized so we rescale to obtain the renormalized frequency *F* (*j, k*), e.g.

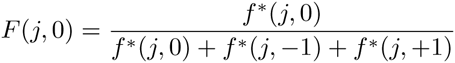

which will be used to compute the expansion time for marker *j*. There are similar formulae if the singularity was at *k* = −1. This is illustrated in figure 3.

However there is sampling error both in the frequencies and the *μ*_*j,*1_, μ_*j,-*1_. So we bootstrap taking into account these uncertainties, running the computation thousands of times. Generally we find the branch singularity is always one of *k* = 0, +1, *-*1 with no SD. In a few cases the singularity may seem to wander between *k* = 0, +1, *-*1. So in the case of a wandering singularity we obtain a distribution over *k* = 0, +1, *-*1 with a mean and SD. In these cases we find the singularity is relatively small and does not make much difference to the final result. However to have a stable method we do not throw out these wandering singularities but in the algorithm use the mean to average between *k* = 0 and *k* = *±*1, e.g. if the mean is *k* = 0 then we use the original unreduced frequency.

Notice that we assume at most one side branch. In theory there could be many and solving for these produce even better approximations to the present data. In fact you could get perfect matching but find the atoms were created yesterday! The thing is that while many markers show significant deviation from a uniform flow from a single patriarch, after we have carried out reduction for one possible side branch we find no significant difference from a uniform flow, i.e. the difference is within the SD. This is of course an approximation, the next level beyond Zuckerkandl and Pauling, but given the noise in the data perhaps the best we can do. Later we further reduce the effect of outliers by using robust statistics.

Reducing the singular lineages increases the frequency *f* (*j,* 0) of the mode and decreases the computed *TMRCA*. But as the method of reducing singularities does not respect higher frequencies *f* (*j, k*) it follows the KAPZ formula cannot be used and instead we use the probability of no mutations, i.e. solve

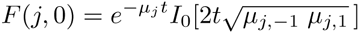

This is done for each DYS marker *j*, giving expansion times *t*_1_,…*t*_*N*_ for each marker, with computed CI. (An extra fixed source of error is the uncertainty in the mutation rates which we deal with later). We find the reduction of singularities makes striking difference to the *t*_*j*_ of the effected markers, often a reduction of *∼* 50% for *TMRCA*.

Now the existence of side branches implies that the main branch could itself have been the side branch for an earlier branch that did not survive. Thus we do not expect the expansion times *t*_1_,…*t*_*N*_ for each marker to be essentially equal., i.e they are not within the SD of each other. Indeed we see that the distribution of the times *t*_*j*_ for different markers are almost certainly not randomly arranged about a single *TRMCA T* but distributed from *T* to the present. This is seen whether you use reduction or not, or our mutation rates or not. (For a given population one could scale mutation rates to get equal *t*_*j*_, but then applying these adhoc mutation rates to other populations does not yield the same values). The spread out distribution of surviving branches is another verification of our theory of many extinctions, few survivors. The distribution of the times *t*_*j*_ for different markers we call the branching distribution, which is now discussed.

### The Branching Distribution

The times *t*_*j*_ for different markers are sorted from the youngest to the oldest, forming a sequence 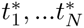. The generation of these branches is by an unknown probability distribution *dτ*_0_ over [0, *T*]. We model *dτ*_0_ by assuming a surviving lineage is generated at random with probability *β*Δ*t* in time period [*t, t* + Δ*t*], multiplied by the probability that the branching hasn’t already occurred. The constant *β* averages fertility and extinction rates, the chance of a new lineage surviving. As *β → ∞* we get current theory where all lineages originate from a single patriarch at time *T*. Simulations with the data show that *β* varies in the range 1 to *∞*. We make no a priori estimate of *β*, unlike Bayesian methods where an overall fertility rate is a predetermined parameter. Instead our stochastic simulation will find the most likely *β, T* in each case. Assuming independence, then the generation of branches follows the well known exponential distribution:

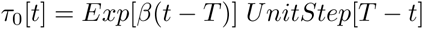

Notice this implies a finite probability that some markers have essentially zero mutations. This is actually seen in examples. Both the Hamilton Gp A and Macdonald Gp A have number of individuals *n >* 100. For the time scale of *>* 700 years we do not expect there is more than one marker out of 33 which shows absolutely no mutations from the mode. In fact in both cases there are 8 markers where all *n* individuals have exactly the same STR value.

Estimating the parameter *T* for an exponential distribution is a well known problem of statistics. Kendall proved the best estimate for *T* would be max *t*_*j*_. Unfortunately there is also considerable error *λ*_*j*_% for the mutation rates *μ*_*j*_. Later we give a method for reducing this error, even so we find the SD in the range 10% - 30% which gives corresponding range in error for each *t*_*j*_. We understand that the *t*_*j*_ are being generated by the distribution *dτ*_0_ but superimposed on this is a further uncertainty due to mutation rates etc. In particular the largest *t*_*j*_ may be wildly inaccurate. Also we found that simply taking the average consistently underestimates the *TMRCA* by a wide margin.

Assuming the mutation rates have normal distribution with mean *μ*_*j*_ and variance 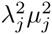, the *t*_*j*_ have SD *t*_*j*_*λ*_*j*_. Thus the actual data for 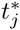 has probability density function for *s >* 0

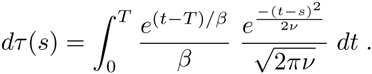

The variance *ν* depends on two sources. First from the uncertainty in mutation rates, for each marker we get variance 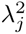, giving total

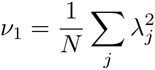

However a small sample also has inherent error from sampling. We are measuring the probability that there is a mutation. This is binomial with probability

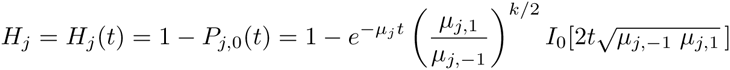

Hence for sample size *n* there is variance *H*_*j*_(1 - *H*_*j*_)*/n*, so the variance in time due to this is scaled by the derivative giving:

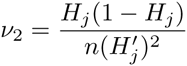

The function 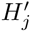 has actually to be computed as an inverse function depending on *H*_*j*_. Therefore the total variance averaged over all *N* markers is *ν* = *ν*_1_ + *ν*_2_. Although for large samples (*n >* 1000) the second term is insignificant it does effect the results once you get to *n* = 100. In our algorithm the branching distribution is used to generate large numbers of random branching times so as to bootstrap error estimates. In turns out much faster to compile the distribution function as a table which can be repeatedly called on.

### Robust Statistics: Estimating *T*

Inaccurate large values of 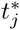 are mitigated by using “robust” statistics with quintiles instead of means/variances. Using FTDNA data we began with 37 markers. However the 4 markers of DYS464 are unordered and cannot be used. Also we find that markers DYS 19/394, 385b, 459b, CDYb have errors *>* 33% in mutation rates so are not used. (These are some of the most popular ones in the literature!). So usually we have *N* = 29 markers and take “quintiles” 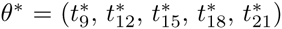. This means that tail end data is not discarded but kept as the information there are 8 values of 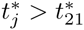, which effectively deals with outliers. Bootstrap methods give the confidence interval CI for each quintile.

Thus we wish to find the best estimate of *T* given *θ*\* (and CI). This well known statistical problem was investigated by Stochastic Simulations (SS). We also tried Maximum Likehood Methods which gave similar results but with larger CI. Monte-Carlo Methods are used to produce very large numbers (*∼*10^7^) of *T, β* with corresponding Distribution. These randomly generate ordered times (*s*_1_*…s*_29_) for which we take the quintiles *θ* = (*s*_9_, *s*_12_, *s*_15_, *s*_18_, *s*_21_). We filter by requiring that *θ* close to the data *θ*\*, i.e. ∥θ* - θ ∥ < 2*SD*. This gives a stochastic neighborhood *U* of θ* typically containing *>* 10^5^ sets of data but with *T* is known for each *θ ∈ U*. Thus we can construct a quasilinear estimator:

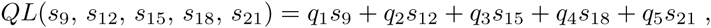

and use least squares over *U* to find constants (*q*_1_, *q*_2_, *q*_3_, *q*_4_, *q*_5_) minimizing

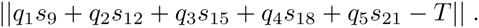

The (*q*_1_, *q*_2_, *q*_3_, *q*_4_, *q*_5_) are computed in MATHEMATICA. We then test them by applying the QL to all of *U*, unsurprisingly

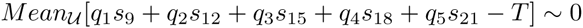

What is important is that we find the uncertainty in the SS itself. Actually this depends on the data and is calculated in each case but for our examples we find

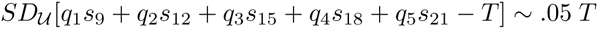

Finally we apply the quasilinear estimator to the experimental data

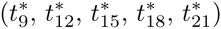

to obtain our best estimate of *T*. Application of *QL* computes the SD for our data, giving part of the overall SD. This must be combined with the SD coming from the uncertainty in the SS. Overall we find that our method has SD *∼*10%, this includes variances from our data, mutation rates and uncertainty in the SS. We also tested with 15 and 7 markers. Here one must use “quintiles” 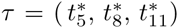,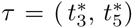 respectively with all the loss of accuracy that implies. See Table 6, 7 for comparisons using 29, 15, 7 markers on same data.

**Figure 8.**
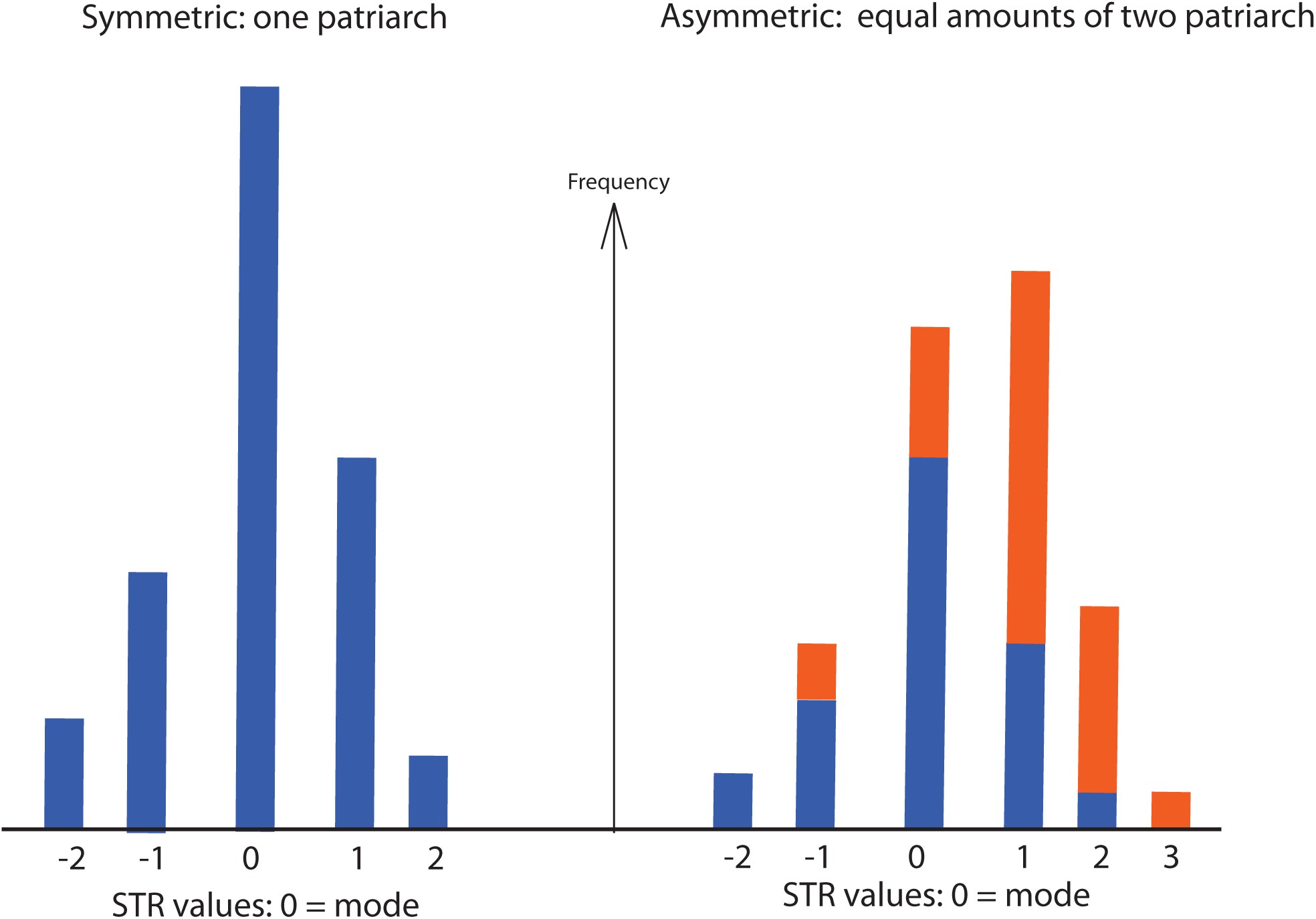

## Supplementary Material 2: Accurate Mutation rates

Any genetic clock depends on reasonably accurate mutation rates. The mitosis method looks for mutations in sperm samples. Forensics uses father-son studies. However typical rates of *μ* = .002 would require nearly 50, 000 pairs to get an SD of 10%. Small samples have meant large errors. The pedigree approach is to study large family groups with well developed DNA/genealogy data. So inverting the KAPZ formula would yield accurate rates. However, *singular lineages* makes this problematic. Genealogical data might give mutation rates much greater than the biochemical rates because kin selection etc tend to exaggerate the apparent mutation rate. An inspection of 10 different sources finds mutation rates claiming SD *∼*10% yet they differ from each other by up to 100%. We describe a new method.

To compute our rates we apply our theory to the large DNA projects for the SNP M222, L21, P312, U106, R1b1a2, I1, R1a1a. This avoids dealing with populations such as family DNA projects which are self selecting, i.e only those with the correct surname which neglects distant branches. Also we have very large samples, our average *n >* 1000. Greater accuracy should come from more generations and individuals. The problem is that we do not know their *TMRCA*.

### Asymmetric Mutation

However before computing mutation rates we must consider asymmetric mutations, i.e. the left and right mutation rates *μ*_*j,-*1_ ≠ μ_*j,*1_. For a uniform stochastic process we again use the asymmetric ratio

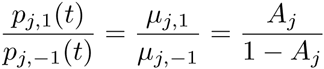

to define the *asymmetric constant A*_*j*_ ∈ [0, 1] for marker *j*. For example *A*_*j*_ = 0.5 is complete symmetry. Of course singularities will effect this ratio, however these only occur *<* 50% of markers. Thus for each marker, SNP we compute this ratio. We find the SD for each SNP is relatively small while the difference between SNP can be large. However for each marker, using 8 SNP enables outliers to be easily removed leaving allowing us to use simple linear regression: i.e. average of the *A*_*j*_ over the remaining SNP groups. We see that asymmetry is a real effect: 50% of the *A*_*j*_ are more than two SD from symmetry *A*_*j*_ = 0.5.

Observe this is significant. The total second moment is

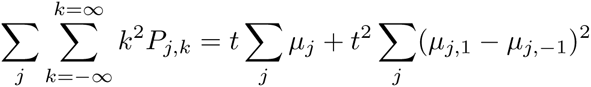

So using all our 33 DYS markers with our *μ*_*j*_, we compute constants

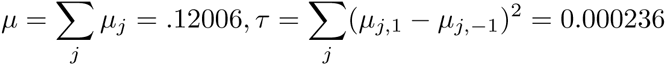

The KAPZ formula gives variance *V* = *μt* compared to the corrected formula *μt* + *τt*^2^. The uncorrected KAPZ gives an overestimate *>* 400% for *>* 200 generations. This effect can be nullified by using the mean instead of the mode, variance instead of the second moment, however failing to do so gives a large error. Furthermore other methods which assume symmetric mutations will also be inaccurate. Having estimates on the asymmetry is essential to our method because we find singular lineages by looking for asymmetry in the data. Any such anomaly needs to be significantly greater than the natural asymmetry.

### Mutation Rates as a fixed Point

Next we compute mutation rates using 8 very large SNP groups. First, using the asymmetric constants we find singular lineages and reduce their effect. We take account of the error in the *A*_*j*_ by a bootstrap technique, which gives the variance for each frequency *f* (*j,* 0). For a given SNP *k* if markers *j* started their expansion at the same time TMRCA *T*_*k*_ we could calculate mutation rates *μ*_*j*_ via

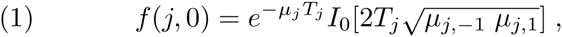

or rather average the 8 different *μ*_*j*_ we would obtain. However because of branching caused by extinction of lineages the different markers do not originate at the same time but at different times *t*_*j*_. In this case we expect these *t*_*j*_ to be randomly distributed about the log mean over a middle set of times *t*_*j*_. So, for each SNP group *k* = 1, *‥*8 define mean time *T*_*k*_, not the TMRCA but the mean log mean over a middle set of markers, which is less. We find that this is very stable. So for a fixed marker *j* the data *τ*_*k,j*_ = *t*_*j*_ - *T*_*k*_ should be randomly distributed about zero over the different SNP *k* = 1, *‥,* 8. However the wrong choose of *μ*_*j*_ would give a bias. In fact this is what we see if the mutation rates *μ*_*j*_ = .002 were chosen. In appendix graphs show the *τ*_*k,j*_, *k* = 1, *‥*8 bunched around a nonzero point. Thus we try to find *μ*_*j*_ so that the *τ*_*k,j*_, *k* = 1, 2, *‥*8 has mean zero. However the *τ*_*k,j*_, *k* = 1, 2, *‥*8 depend nonlinearly on the rates *μ*_*j*_, as does the mean *T*_*k*_, *k* = 1, *‥*8. We find this nonlinear regression problem is solved by an iterative scheme which starts with any reasonable set of DNA rates, finding any reasonable choice iterates to the same final answer. So choose *μ*_*j*_ = .002 to begin. Suppose at some stage we have apparent mutation rates *μ*_*j*_. Then, for each SNP, and each marker we solve equation (1) to obtain the apparent *t*_*j*_. For each SNP *k* = 1, *‥*8 we compute the mean log time *T*_*k*_. At the next step we get new rates 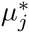 from

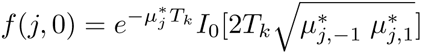

Averaging 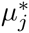, *k* = 1, *‥*8 we get our next set of *μ*_*j*_ of mutation rates. However this method would be effected by a marker showing a singular lineage. Fortunately these are few in number and by comparison between the different SNP we remove the outliers. We then repeat the process, computing *T*_*k*_ again with the new rates, and another set of mutation rates. So we have an iterative process.

One problem is that the iterates could tend to decrease to zero or increase to *∞*, as we are only calculatin),g relative rates. To prevent this we renormalize after each iteration so the total Σ*μ*_*j*_ is constant. We found the iterative scheme quickly converges to a fixed set of mutation rates, unique up to a constant factor. The CI is computed by bootstrap parametrized by the uncertainties in data and the asymmetric constants. In figure we show the distribution of *τ*_*k,*1_, *k* = 1.2, *‥*8 before and after the first iteration.

### The generation factor *γ*

This method does not give absolute mutation rates but *relative* mutation rates *μ*_*j*_*γ*, where *γ* is universal time scale constant. To find *γ* we apply our method to compute the *T* = *T M RCA* of three famous DNA projects and choose *γ* so the scaled *T /γ* best fits the historical record. We choose the DNA projects for the O’Niall(M222), Gp A of Macdonald (R1a1a) and Gp A of the Hamiltons (I1). These are large groups with characteristic DNA and fairly accurate times of origin. Of course finding one constant *γ* from three projects is inherently more accurate than using one project to find 33 different mutation rates. Actually assuming a generation of 27*years* these three projects yield *γ* = 1 with about 5% error, i.e. there is no actual need for this correction. This is a constant error (like uncalibrated ^14^*C* dating).

Thus *γ* is related to the length of a generation. Most researchers use 25*yrs* for *t >* 500*ybp* and 27*yrs* for *t <* 500*ybp*. Balaresque and al used 30*yrs* based on Finer who sees a 30*yr* generation for modern hunter-gatherers. (Although for most of the time R1b1a2 were subsistence farmers and not hunter gatherers.) At first glance our theory allows any nominal generation as it really doesn’t matter, being included in the *γ* factor which we compute in years not generations. Actually its not as simple as that. While our three DNA projects being post 1000AD elites have a 27*yr* generation the problem is what to do for *t >* 2000*ybp*. Now 25*y* may be appropriate for subsistence farmers but we found that singular lineages of the elite have exaggerated effect so 27 years seems appropriate.

Mutation Rates: Hamilton vs Mitosis and pedigree

| # | DYS | Hamilton | DH-SD | DH+SD | Burgella<br>(mitosis) | B(m) - SD | B(m) +SD | Burgella<br>(regress.) | B(r) - SD | B(r)+SD | NIST | FTDNA |
| --- | --- | --- | --- | --- | --- | --- | --- | --- | --- | --- | --- | --- |
| 1 | 393 | 0.72 | 0.56 | 0.92 | 1.03 | 0.60 | 1.77 | 2.60 | 2.16 | 3.119 | 0.08 | 1.43 |
| 2 | 390 | 2.52 | 1.87 | 3.40 | 2.12 | 1.49 | 3.03 | 4.66 | 3.92 | 5.53 | 2.40 | 5.32 |
| 3 | 19/394* | 1.30 | 0.64 | 2.64 | 2.19 | 1.55 | 3.09 | 2.84 | 2.53 | 3.182 | 2.38 | 1.45 |
| 4 | 391 | 4.98 | 3.56 | 6.96 | 2.72 | 1.98 | 3.72 | 2.02 | 1.74 | 2.343 | 2.88 | 4.15 |
| 5 | 385a | 1.26 | 1.00 | 1.58 |  |  |  |  |  |  | 2.10 | 5.68 |
| 6 | 385b* | 3.13 | 1.87 | 5.25 |  |  |  |  |  |  | 2.10 | 5.68 |
| 7 | 426 | 0.07 | 0.04 | 0.11 |  |  |  | 0.46 | 0.25 | 0.842 |  | 0.26 |
| 8 | 388 | 0.22 | 0.15 | 0.32 | 0.42 | 0.02 | 2.36 | 0.46 | 0.25 | 0.843 |  | 0.25 |
| 9 | 439 | 3.76 | 3.07 | 4.60 | 5.48 | 4.17 | 7.19 | 2.90 | 2.58 | 3.248 |  | 4.95 |
| 10 | 389-I | 1.93 | 1.61 | 2.31 | 2.53 | 1.79 | 3.57 | 2.20 | 1.92 | 2.517 | 1.88 | 2.23 |
| 11 | 392 | 0.36 | 0.23 | 0.56 | 0.43 | 0.20 | 0.94 | 0.48 | 0.26 | 0.861 | 0.58 | 1.59 |
| 12 | 389b | 2.96 | 2.42 | 3.62 | 3.17 | 2.33 | 4.31 | 2.54 | 2.26 | 2.867 | 2.96 | 2.72 |
| 13 | 458 | 7.99 | 6.83 | 9.35 | 6.88 | 5.16 | 9.17 | 4.78 | 3.91 | 5.838 | 10.80 | 6.30 |
| 14 | 459a | 0.39 | 0.29 | 0.53 |  |  |  |  |  |  |  |  |
| 15 | 459b* | 2.98 | 1.54 | 5.76 |  |  |  |  |  |  |  |  |
| 16 | 455 | 0.16 | 0.12 | 0.22 |  |  |  | 2.14 | 1.74 | 2.63 |  | 0.46 |
| 17 | 454 | 0.11 | 0.08 | 0.15 |  |  |  | 2.18 | 1.78 | 2.674 |  | 0.47 |
| 18 | 447 | 3.80 | 2.93 | 4.92 | 4.56 | 1.55 | 13.32 | 0.74 | 0.37 | 1.467 |  | 4.00 |
| 19 | 437 | 0.99 | 0.73 | 1.35 |  |  |  |  |  |  | 1.50 | 2.15 |
| 20 | 448 | 1.16 | 0.81 | 1.66 |  |  |  |  |  |  | 1.80 | 2.71 |
| 21 | 449 | 11.70 | 9.16 | 14.94 | 18.97 | 9.22 | 38.63 | 9.64 | 6.85 | 13.56 |  | 7.84 |
| 22 | 460 | 2.63 | 2.08 | 3.33 | 3.82 | 1.63 | 8.92 | 2.49 | 2.06 | 3 |  |  |
| 23 | GATAH4 | 3.93 | 3.28 | 4.71 | 2.43 | 1.80 | 4.211 | 2.19 | 1.91 | 2.515 | 2.51 |  |
| 24 | YCA IIa | 0.32 | 0.22 | 0.46 |  |  |  |  |  |  |  |  |
| 25 | YCA IIb | 1.40 | 1.03 | 1.90 |  |  |  |  |  |  |  |  |
| 26 | 456 | 8.10 | 5.56 | 11.80 | 4.50 | 3.16 | 6.42 | 3.27 | 2.75 | 3.881 |  |  |
| 27 | 607 | 2.15 | 1.72 | 2.69 |  |  |  | 3.73 | 3.27 | 4.268 |  | 4.10 |
| 28 | 576 | 10.65 | 8.72 | 13.01 | 16.22 | 8.55 | 30.53 | 4.18 | 3.48 | 5.034 |  | 10.20 |
| 29 | 570 | 4.60 | 3.27 | 6.48 | 12.61 | 6.12 | 25.80 | 4.20 | 3.49 | 5.059 |  | 7.90 |
| 30 | CDYa | 14.71 | 12.37 | 17.49 |  |  |  |  |  |  |  | 35.30 |
| 31 | CDYb* | 13.40 | 2.68 | 67.00 |  |  |  |  |  |  |  | 35.30 |
| 32 | 442 | 2.90 | 2.38 | 3.54 |  |  |  | 1.93 | 1.64 | 2.256 |  |  |
| 33 | 438 | 0.43 | 0.34 | 0.55 |  |  |  |  |  |  | 0.70 |  |

Mutation rates $10^{-3}$ .
Comparison with Mitosis rates: Burgella summarized 20 different Mitosis sets, e.g. NIST (shown), then used averages(m) and regression(r)
FTDNA uses pedigree studies
\*Too inaccurate to use

Asymmetric rates

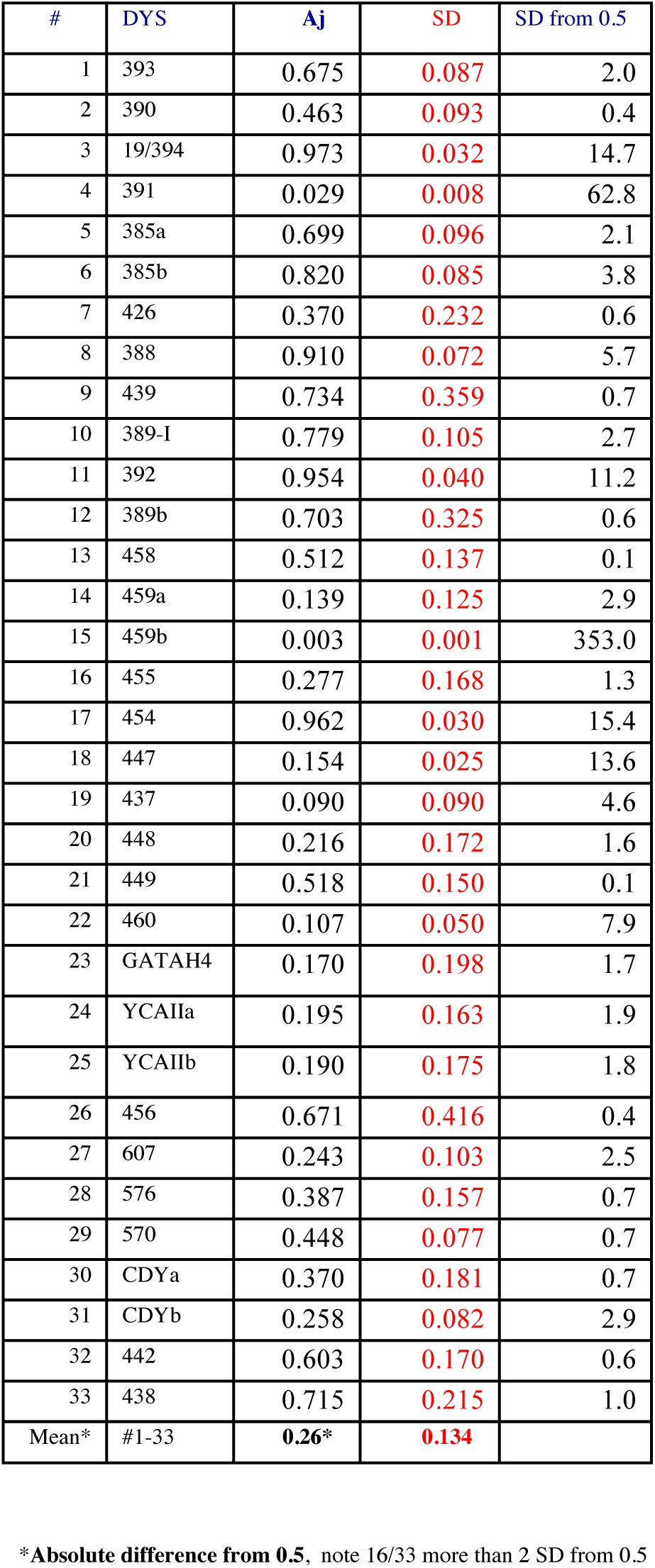

The log distribution of *τ*_k,1_, *k* = 1.2, *‥*8 before iteration at marker *j* = 1, ie DYS 393, but after reduction ^1^(*μ*_*j*_ = .002). The SNP are colored:

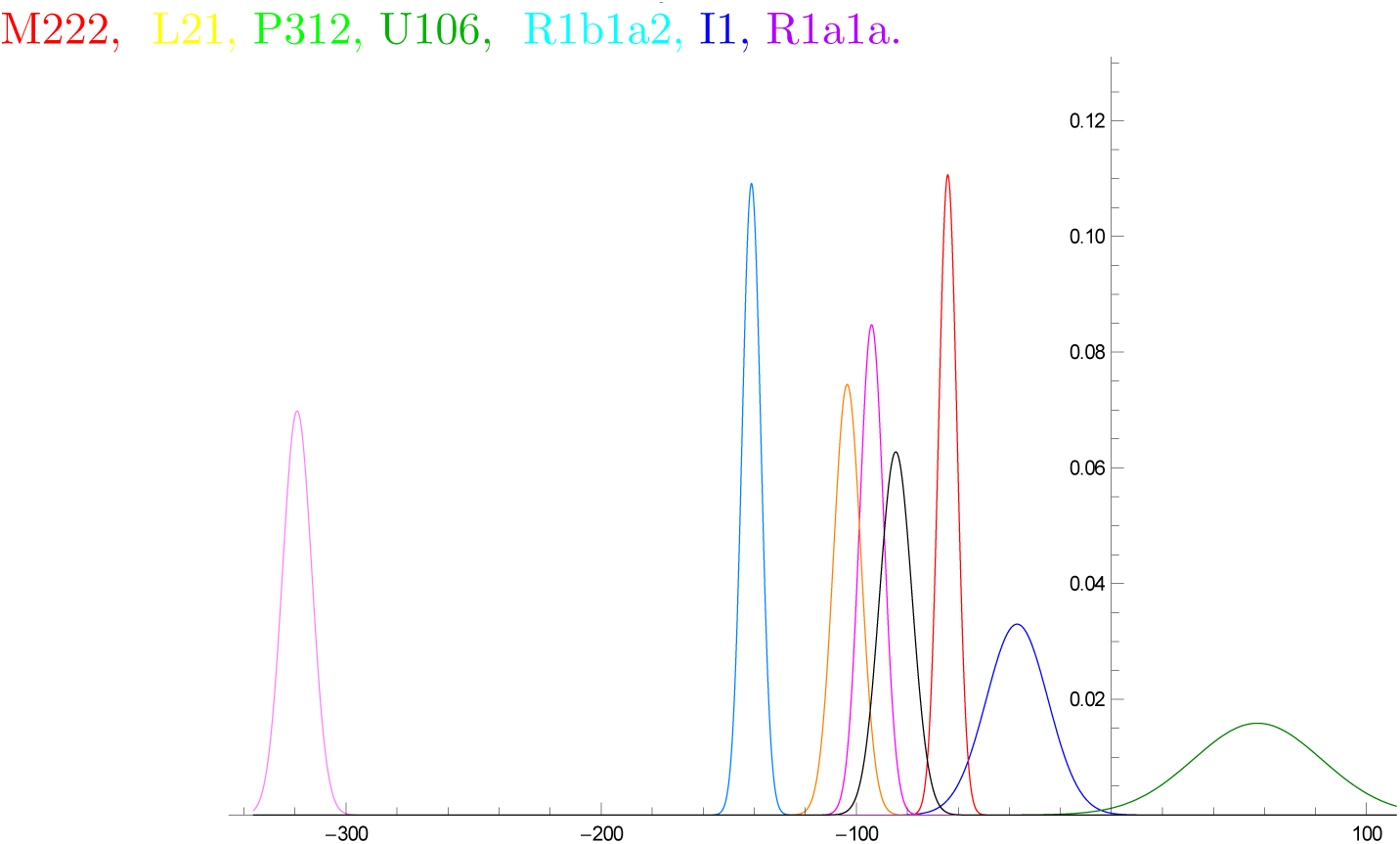

After just one iterate we get

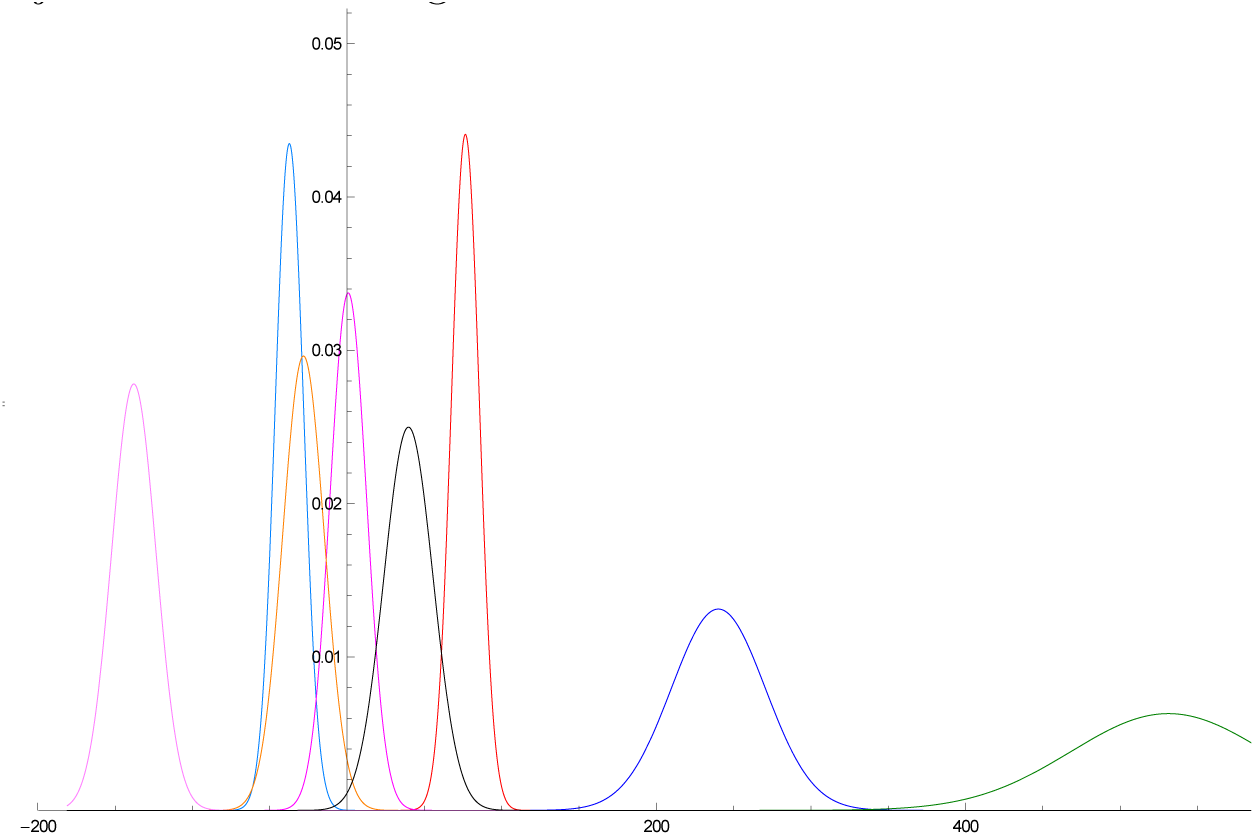

So 5 of our *τ*_*k,*1_, *k* = 1.2, *‥*8 bunch around zero, outliers are U106 and I1.

The iterative scheme converges to stable values very fast, 7 iterates is enough.

## Supplementary Material 3: Reduction of Singular Lineages vs KAPZ

We compare results for our method with KAPZ, for the same data, 29 markers and our mutation rates

First we compare for groups with medieval expansions

| *Group* | *(n)* | *RSL* | *[SD]* | *KAPZ* | *[SD]* | First Known |
| --- | --- | --- | --- | --- | --- | --- |
| *Hamilton* | (144) | 1358AD | [140] | 1130AD | [255] | W. Hamilton 1300AD |
| *Macdonald* | (95) | 900AD | [250] | 530AD | [566] | Somerled 1100AD |
| *O’Niall* | (713) | 200AD | [225] | 20BC | [364] | Niall 300AD |

Next we compare SNP G2a2b, R1b1a2, R1a1a, I1, L21, U106, J2, P312:

| *SNP* | *(n)* | *RSL* | *[SD]* | *KAPZ* | *[SD]* |
| --- | --- | --- | --- | --- | --- |
| *G2a2b* | (1221) | 5359BC | [900] | 4840BC | [1257] |
| *R1b1a2* | (460) | 3700BC | [625] | 5490BC | [2144] |
| *R1a1a* | (1270) | 3800BC | [700] | 3670BC | [1066] |
| *I1* | (2898) | 1800BC | [400] | 2400BC | [1061] |
| *L21* | (1029) | 1600BC | [325] | 3270BC | [1063] |
| *U106* | (1533) | 2400BC | [440] | 2530BC | [628] |
| *J2* | (1241) | 15500BC | [2600] | 11700BC | [2990] |
| *P312* | (971) | 2240BC | [420] | 2900BC | [632] |

RSL and KAPZ will give similar results if there is a fast expansion and thus insignificant singular lineages and branching. Actually this is to be expected sometimes, i.e. it is not surprising that the results using RSL and KAPZ for O’Niall, R1a1a, U106 are very similar.

However in other cases the KAPZ results are about 30% too old. In the case of the Hamiltons and Macdonalds absurdly so. For R1b1a2 it gives an early Neolithic age, compared with eneolithic for R1a1a, yet these have been dated to the same Yamanya times. The KAPZ dates for L21 “Celtic” is nearly 2000 years before Urnfelder Culture.

Of course one might try to “improve” KAPZ by increasing the mutation rates by 33% so the KAPZ times are decreased by 25%. Then the medieval dates look reasonable but we find 3100BC for G2a2b which is too late. For R1a1a we would get 2300BC which is not only too late but significantly different from the 3600BC for R1b1a2. Also G2a2b would be predated by R1b1a2 even though the latter has never been found in Neolithic sites of Europe. Getting consistent results across the span of history was a problem of previous clocks.

## Footnotes

1 The calculations and figures for all 33 markers is shown in SM

